# Non-Gaussian, transiently anomalous and ergodic self-diffusion of flexible dumbbells in crowded two-dimensional environments: coupled translational and rotational motions

**DOI:** 10.1101/2021.09.06.459157

**Authors:** Kolja Klett, Andrey G. Cherstvy, Jaeoh Shin, Igor M. Sokolov, Ralf Metzler

## Abstract

We employ Langevin-dynamics simulations to unveil non-Brownian and non-Gaussian center-of-mass self-diffusion of massive flexible dumbbell-shaped particles in crowded two-dimensional solutions. We also study the intra-dumbbell dynamics due to the relative motion of the two constituent elastically-coupled disks. Our main focus is on effects of the crowding fraction *ϕ* and the particle structure on the diffusion characteristics. We evaluate the time-averaged mean-squared displacement (TAMSD), the displacement probability-density function (PDF) and the displacement autocorrelation function (ACF) of the dimers. For the TAMSD at highly crowded conditions of dumbbells, e.g., we observe a transition from the short-time ballistic behavior, via an intermediate subdiffusive regime, to long-time Brownian-like spreading dynamics. The crowded system of dimers exhibits two distinct diffusion regimes distinguished by the scaling exponent of the TAMSD, the dependence of the diffusivity on *ϕ*, and the features of the displacement-ACF. We attribute these regimes to a crowding-induced transition from a viscous to a viscoelastic diffusion medium upon growing *ϕ*. We also analyze the relative motion in the dimers, finding that larger *ϕ* suppress their vibrations and yield strongly non-Gaussian PDFs of rotational displacements. For the diffusion coefficients *D*(*ϕ*) of translational and rotational motion of the dumbbells an exponential decay with *ϕ* for weak and a power-law *D*(*ϕ*) ∝ (*ϕ* – *ϕ*⋆)^2.4^ for strong crowding is found. A comparison of simulation results with theoretical predictions for *D*(*ϕ*) is discussed and some relevant experimental systems are overviewed.

## I. INTRODUCTION

### A. Historical prologue

In 1827, R. Brown observed under a microscope the erratic motion of micron-sized granules released from pollen grains^1^. The paradigmatic stochastic process— named later Brownian motion (BM)—was physically interpreted by A. Einstein^2^ in his *annus-mirabilis* 1905 paper (and in later studies^3,4^). The probability-density function (PDF) *P*(*x, t*) of one-dimensional BM with the long-time diffusion coefficient *D* satisfies the diffusion equation, 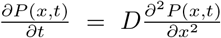, introduced in 1855 by A. Fick^5^. The derivation of BM^2^ is based on the assumptions of independence of motion of a given particle from other particles and of identically distributed particle displacements that become independent after a finite correlation time (yielding a finite second moment). The era of “random walks” started with the papers of L. Bachelier^6^, W. Sutherland^7^, A. Einstein^2^, K. Pearson^8^, and M. Smoluchowski^9^.

The solution of the diffusion equation for a single particle at position x with P. Dirac’s^10^ delta-function initial condition *P*(*x, t*_0_ = 0) = *δ*(*x* – *x*_0_) is the Gaussian PDF [after C. Gauß^11^] at time *t*,

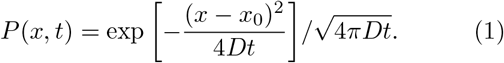

This PDF yields the linear growth of the mean-squared displacement (MSD),

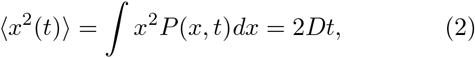

a hallmark of BM. All ensemble-averaged quantities are denoted by angular brackets below. The diffusivity of a *spherical* tracer of radius *R* at the absolute temperature 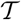, denoted 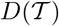, is linked to the (dynamical) fluid viscosity *η* via the Einstein-Smoluchowski-Stokes relation^2,9^,

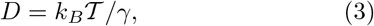

where *γ* = 6*πηR* is the drag constant and *k_B_* is the Boltzmann constant.

In 1908, P. Langevin derived Eqs. (2) and (3) using a microscopic inherently stochastic approach^12^ for a spherical particle of finite mass *m*. Starting from I. Newton’s second law^13^—from his *annus mirabilis* 1666 (marked also by the “Great Plague”)—using G. Stokes’ drag— introduced in 1851^14^ for slowly moving bodies in viscous fluids—and the random force *ξ*(*t*) [taken to be centered white Gaussian noise], the *stochastic* differential equation was postulated,

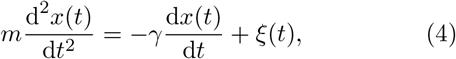

(named later after Langevin). As Eq. (4) takes inertia into account, at short times when *t* ≪ *τ*_0_, where

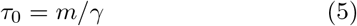

is the characteristic time of momentum relaxation for translational motion [the damping constant is 1 /*τ*_0_], a ballistic behavior of the MSD,

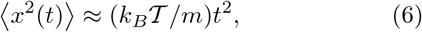

emerges, while the linear MSD growth (2) sets in at long times, for *t* ≫ *τ*_0_^15–17^. The features of BM executed by *ellipsoidal* particles was first theoretically considered by F. Perrin in Refs,^18,19^.

### B. Anomalous diffusion, macromolecular crowding, and spreading of non-spherical particles

Standard BM has been generalized in numerous mathematical models of anomalous diffusion over the last decades^20–24^. These models feature a characteristic nonlinear growth of the MSD with time^24–29^,

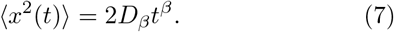

Here, the generalized diffusion coefficient is *D_β_* and the anomalous-diffusion scaling exponent is *β*. The dispersion law for BM follows from (7) for *β* = 1, while subdiffusion corresponds to 0 < *β* < 1, superdiffusion to *β* > 1 (for actively driven systems), and hyperdiffusion— often found in highly nonequilibrium and dynamically “accelerating” systems—to *β* > 2. Following growing evidence for experimental systems, a number of new statistical models were proposed in recent years to describe anomalous diffusion^24–27^. The list includes fractional BM^30–34^, generalized/fractional Langevin equation motion^35–37^, scaled BM^38–41^, extensions of continuous-time random-walk models^39,42–48^, and heterogeneous diffusion processes^33,49,50^.*

In the cell cytoplasm, the average degree of crowding is the volume fraction taken from the aqueous phase^27,53–71^ by proteins, diverse macromolecules, complexes, and cell organelles. In cells, the fraction of crowders *ϕ* can reach *ϕ* ~ 30 … 40% with concentrations of various proteins and macromolecules up to ~300… 400 mg/ml. In colloidal systems, the crowding fractions can be even higher triggering liquid-glass phase transitions^72–76^ (see Refs.^77,78^ for glass transitions in 2D).

Space-filling and crowding in excluded-volume solutions—causing effective “labyrinthization” and “ porosityzation” of the diffusive environment (also with tracer-crowders interactions^79^)—is considered as a cause of subdiffusion of passive particles in densely crowded systems. This is interlinked to implications of viscoelasticity in space-restricted surroundings and in the presence of multiple boundaries, nontrivially affecting the tracer diffusion, even in the limit of transport with small diffusion coefficients and particle speeds.

Normal and anomalous diffusion in the crowded solutions of anisotropic and/or non-spherical—as well as of spherical but non-isotropically interacting^80,81^—molecules was examined in a number of studies^82–103^, including some recent experimental investigations^96,99,104^. The diffusive properties of molecules of dumbbell-, ellipsoid-, and rod-like shape^82,96,97,100,101,104–109^, protein molecules^110,111^, active and passive dimers with anisotropic mobility^89,102,106,112–114^, self-propelled massive particles with time-dependent mass^109^, colloidal dimers and flexible multimers^115,116^, stiff self-propelled filaments in crowded solutions^117^, trimer-like tracers in 2D^93,95^ and tetrahedral patchy particles in 3D^81,118^, as well as the dynamics of elongated particles with fluctuating and oscillating shapes^119–121^ were studied recently as well.

A dumbbell-shaped dimer, see Fig. 1A, is an example of a simple linear crowder for which the effects of interparticle interactions and shape variations on the translational and rotational diffusion dynamics can be examined. Under conditions of dense packing, the motion of a given crowder is no longer independent of that of the neighboring crowders, see Fig. 1C. This invalidates one of the assumptions for BM. As a result, subdiffusion can emerge (as we show below) due, in part, to anticorrelations of consecutive displacements of the particles in such a highly crowded system.

**FIG. 1:**
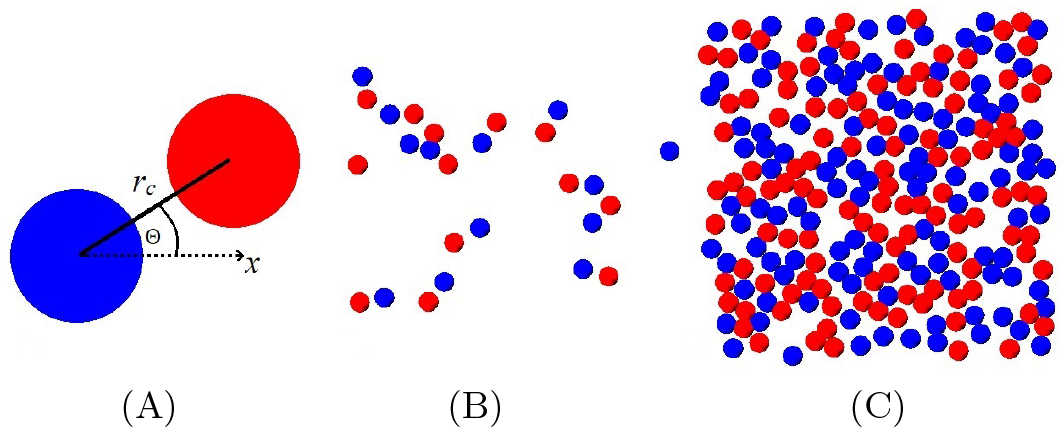
(A) Dumbbell-shaped dimer of our simulation model. (B) Diluted system of dimers at crowding fraction *ϕ* = 0.05. (C) Highly crowded system of dumbbells at *ϕ* = 0.45. We refer the reader to the supplementary video files (with 360 simulation steps corresponding to the duration 0.18 × *δτ* per one second of the videos) illustrating the dynamics of dumbbells at *ϕ* = {0.15, 0.35, 0.65}.

The examples of physical systems where diffusion of anisotropic crowders is of importance include anisotropic crowders in the cell cytoplasm^55,63^, rod-shaped grains^122^ in various granular gases^123–125^, molecular components of ultra-dense ionic liquids^126,127^ and other complex fluids^98,128^, diffusion of proteins and enzymes on/in crowded lipid membranes^29,129–132^ of biological cells, to mention a few.

This paper is organized as follows. We start in Sec. II by describing the model potentials and simulation scheme of dense 2D solutions of dumbbell-shaped crowders. In Sec. III the main results of our computer simulations are presented and examined based on a number of standard statistical quantifiers. The latter include the MSD, the time-averaged MSD (TAMSD), the exponents and transport coefficients, the PDFs, the non-Gaussianity parameter, and various autocorrelation functions (ACFs). In Sec. IV some physical applications are discussed. In Apps. A and B, correspondingly, some details of the simulation scheme and some auxiliary figures are presented.

## II. PHYSICAL MODEL

### A. Interaction potentials and main equations

We consider an ensemble of identical 2D dumbbell-shaped dimers in 2D solution. Each dimer, see Fig. 1A, consists of two disks of diameter *σ* connected by an elastic spring represented by the harmonic potential,

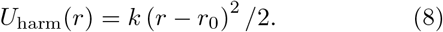

Here, *r*_0_ = 1.5*σ* is the equilibrium disk-to-disk distance and 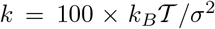 is Hooke’s constant. In addition, the standard Lennard-Jones 6-12 potential is used to parameterize the interactions between the disks,^†^

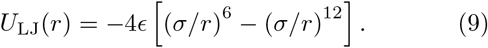

The parameter *ϵ* sets the interaction strength (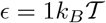 below). The minimum of the interaction energy *U*_min_ is at *r*_max_ = 2^1/6^*σ* ≈ 1.12*σ*. The repulsive branch of the Lennard-Jones potential (9) at *r* < *r*_max_ is used in the simulations, yielding the Weeks-Chandler-Anderson potential^137–139^. It is given by expression (9) shifted up by *U*_min_ at *r* < *r*_max_ and by *U* = 0 at *r* > *r*_max_. As *r*_0_ > *r*_max_ the disks of the neighboring dimers can get closer than those of the same one.

The respective Langevin equation for the position of the ith monomer-disk **r**_*i*_(*t*) is given by

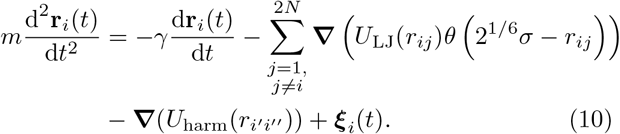

Here, *θ*(*x*) is the unit-step function, the sum runs over all 2*N* disks of *N* dumbbells, and ***ξ***(*t*) is Gaussian white noise with zero mean 〈***ξ***(*t*)〉 = 0 and delta-function^140^ correlations

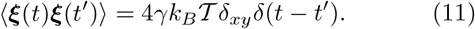

Here, *δ_xy_* is Kronecker’s delta-symbol that renders the noise in (11) independent for *x* and *y* components. The noise strength is coupled to the temperature and the friction coefficient via the fluctuation-dissipation relation. The distance 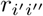 in Eq. (10) is the separation of the two disks *i*′ and *ii*″ in the same ith dimer.

### B. Self-diffusivity and crowding

All dimers are located in a square box of area *L*^2^, but due to periodic boundary conditions their motion is effectively unconfined. For *N* dimers occupying the area *A* = 2*π*(*σ*/2)^2^ the packing fraction *ϕ* in the box is *ϕ* = *NA/L*^2^.^‡^ We fix the system size to *L* = 20*σ* for simulations to be manageable on a PC without causing artificial self-interactions. The number *N* of dimers and the respective *ϕ* fractions used in our simulations as listed in Tab. I. As our main interest is to study the consequences of crowding and thus fluid inertia^145–147^ and hydrodynamic interactions^90,148,149^—often non-negligible for actively-driven and quickly moving particles—are *not* explicitly included in our model of the passively diffusing dimers.

**TABLE I:** Numbers of dimers *N* and the corresponding packing fractions *ϕ* used in simulations.

|  |  |  |  |  |  |  |  |  |
| --- | --- | --- | --- | --- | --- | --- | --- | --- |
| $N =$ | 1 | 13 | 38 | 63 | 89 | 115 | 140 | 166 |
| $\phi \approx$ | 0.004 | 0.05 | 0.15 | 0.25 | 0.35 | 0.45 | 0.55 | 0.65 |

### C. Simulation algorithms and system equilibration

We set *γ* = 1 in most of the computer simulations— at *m* = 1 and *k* = 100 a free isolated dimer has a small damping ratio and, thus, executes underdamped motion—and compute time series of length *T*_sim_ = 10^8^ × Δ*t*. We use the standard combination of parameters^150^ for defining the characteristic time scale^93^,

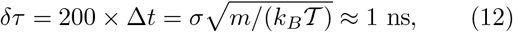

where the disk diameter is set to *σ* = 6 nm, the mass *m* is estimated from the average molecular weight of crowding molecules in the nucleus^151^ (with MW≈ 68 kDa^60^), and the temperature is set to 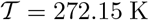. The length of simulated traces is *T*_sim_ = 0.5 ms. The latter is, thus, comparable to typical time-scales in computer-simulations data-sets and single-particle-tracking experiments. All distances and times are expressed below in units *σ* and *δτ*, respectively.

We equilibrate the system for the time *T*_eq_ before the actual measurements start, see Fig. S1 for a detailed scheme. For a dimer of size *R*_cr_ and diffusivity *D*(*ϕ*) we define *T*_cr_ as the typical time for a dimer to diffuse over a distance of its own size^93^, i.e. 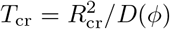. To allow proper mixing of the dimers in the course of simulations, we equilibrate for *T*_eq_ = 2 × 10^4^ × *δτ* for all *ϕ* values (so that *T*_eq_ ≥ 10^2^ × *T*_cr_).

At the start of the simulations, the dimers are placed in a regular pattern and the disks of each dimer are further than *r*_0_ apart. The memory of these initial conditions is lost in the results presented below after equilibration of the system, for all *ϕ* values.^§^ After equilibrating the system for *T*_eq_, at every time-step we record four physical observables for each dimer: the *x* and *y* coordinates of its center of mass (COM), denoted as *x*_COM_(*t*) and *y*_COM_(*t*), the angle Θ(*t*) between its axis and the *x*-axis, see Fig. 1A, and the relative distance between the disks in the dimer, *d*(*t*). Their displacements separated by a lag time Δ are denoted as

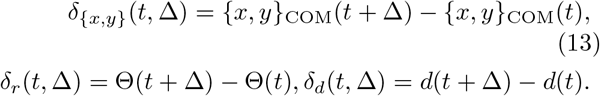

By symmetry, the behaviors of *δ_x_* and *δ_y_* are statistically identical, so we present only the results for the *x*-coordinate below.

We employ the velocity Verlet algorithm^150^ to simulate Eq. (10), see App. A for the detailed description, with the integration time step Δ*t* = 0.05 × *δτ* and the number of iterations 10^8^. We start measuring after the system reaches the equilibrium state. In Fig. S1 we show a simplified flow-chart of simulations: after the initial setup, the integration loop is executed and Eqs. (A5) are repeatedly evaluated.

## III. MAIN RESULTS

Below the data from our computer simulations are analyzed and the diffusive properties of the dumbbell-shaped dimers are examined for both translational and rotational motion. We employ the standard physical observables such as the MSD and the TAMSD, their respective scaling exponents and generalized diffusivities, the displacement PDFs, and the ACFs.

### A. TAMSD and MSD

We compute the translational TAMSD for the COM positions (denoted as 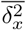) and the rotational TAMSD for the angles Θ (denoted as 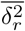) of the *i*th dimer as the sliding averages along respective time series recorded from simulations of Eq. (10) via the standard definition^24,25,93^

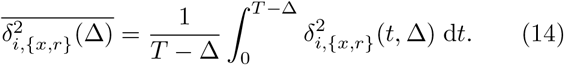

Here, *T* is the total length of the recorded trajectories and Δ is the lag time^24,25^. All time-averaged quantities are denoted by an overline below. The mean TAMSD after averaging over *N* trajectories is

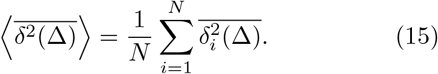

In Fig. 2 we show the behavior of the computed translational and rotational TAMSDs of a single dimer versus the lag time Δ for varying crowding fraction *ϕ*. Initially, for lag times Δ ≲ 0.1 × *δτ*, the diffusion of dimers is found to be ballistic and the magnitude of the mean TAMSDs is almost the same for all *ϕ* fractions examined: in this short-time regime the dimers perform free motion without collisions, with the average speed 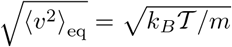 given by Eq. (6). With increasing *ϕ* this regime shrinks to progressively shorter times. The ballistic regime at short times is also detected in simulations of diffusion of a single dimer (corresponding to *ϕ*_min_ = 0.004), with the MSD results presented in Fig. S2 for varying *γ* values.^¶^

**FIG. 2:**
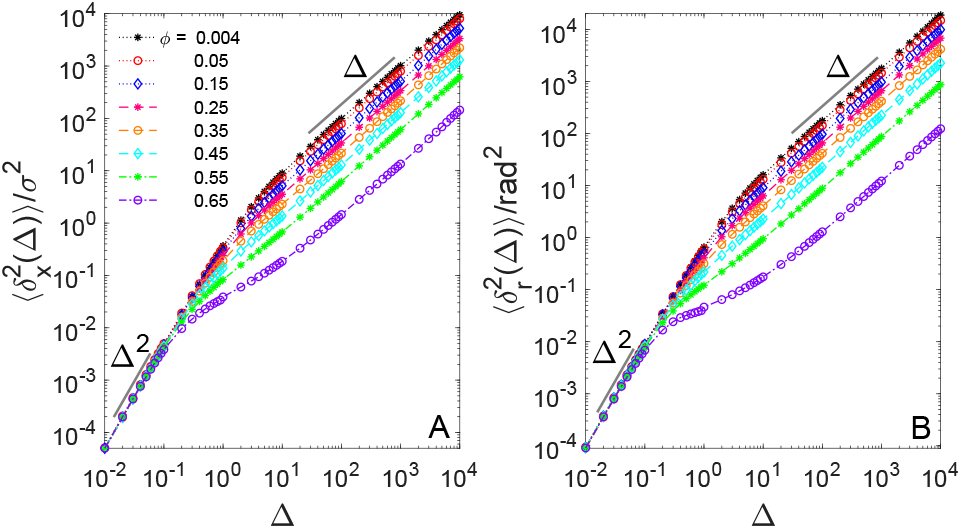
(A) Translational TAMSD in units of *σ*^2^ and (B) rotational TAMSD in units of rad^2^ of the dumbbell-shaped crowders given by Eq. (14) plotted versus lag time for varying crowding fractions *ϕ*. The ballistic and linear asymptotes at short and long times, respectively, are shown in each panel as the thin lines. Parameters: the mass of the disk is *m* = 1, the friction coefficient is *γ* = 1 yielding the velocity-relaxation time *τ*_0_ = 1 × *δτ*, the strength of the potential is 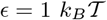, the trace length is *T* = 5 × 10^5^, and the averages are computed for an ensemble of *N* = 40 trajectories. The lag time Δ and the trajectory length *T* are given in units *δτ* ≈ 1 ns in this and all other plots. The values for the crowding fractions *ϕ* are provided in the legend.

Physically, when a dimer diffuses only a fraction of its size from the initial location it does not yet feel any hindrance by collisions with the neighboring crowders and, thus, moves with a constant temperature-dependent velocity, see Eq. (6). This yields the quadratic growth of the MSD [see Eq. (6) and Eq. (16) below] and of the mean TAMSD at short times, as shown in Figs. 2 and S2. As physically expected, as the friction coefficient increases and corresponding the relaxation times for translational and rotational motion become shorter, the region of ballistic MSD growth gets limited to shorter times, see Fig. S2.

At intermediate lag times, a crossover behavior of the TAMSD in the crowded solutions of dimers is observed (see Fig. 2) and the computed mean TAMSDs curves start to *split* for different crowding fractions *ϕ*. This regime is realized in the region of lag times Δ ≈ 0.1… 10 × *δτ*. At the end of this interval, the translational and rotational TAMSDs grow almost linearly with lag time. This ballistic-to-linear TAMSD crossover depends on *ϕ* and is characterized by the first-collision time (see Sec. III B for the analysis of scaling exponents).

Finally, at very long times, both TAMSDs grow linearly with Δ and follow BM (2) with *ϕ*-dependent effective diffusivities (see Sec. III C below).

When comparing the behaviors at *γ* = 1 we observe that for small crowding fractions *ϕ* the MSDs of both translational and angular motions are almost equivalent in the entire range of times to the theoretically expected results for 〈*x*^2^(*t*)〉 and 〈Θ^2^(*t*)〉. Indeed, for the MSD of a single dimer obeying a potential-free version of Eq. (10) one expects the MSD growth^15,16,145,152^

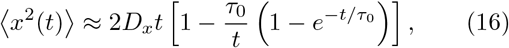

also known in polymer physics for the average extension of a fluctuating worm-like chain (see §127 in Ref.^153^). For the rotational MSD 〈Θ^2^(*t*)〉 we use Eq. (16) with the friction coefficient extracted from the diffusivity results of computer simulations based on relation (3), while the particle mass is now being exchanged with the moment of inertia of the dimer (computed with the potential-equilibrated monomer-monomer separation *r*_0_ = 1.5*σ* using the known mass of the monomer disks). This gives the characteristic time of rotational relaxation of the motion, denoted *τ*_0,Θ_. The diffusion of dimers is anisotropic with respect to their axes at short times, when *t* ≪ *τ*_0,Θ_, and it turns isotropic at long times, when *t* ≫ *τ*_0,Θ_.

The crossover from the short-time ballistic to the longtime linear behavior of the rotational and translational MSD and mean TAMSD takes place at times ~ *τ*_0_ and ~ *τ*_0,Θ_, respectively, for weakly crowded systems and does so monotonically in terms of reduction of the time-local scaling exponent, see Eq. (17) below. In contrast, the crossover emerges at considerably *earlier* times for highly crowded solutions of the dumbbells, see Figs. 2 and 3, accompanied by a *nonmonotonic* variation of the respective exponents *β_x_* and *β_r_*. At high crowding fractions at intermediate times—where the subdiffusive behavior is most pronounced, see also Fig. 3—the theoretically expected and observed MSD are rather disparate in magnitude, see Fig. S3.

**FIG. 3:**
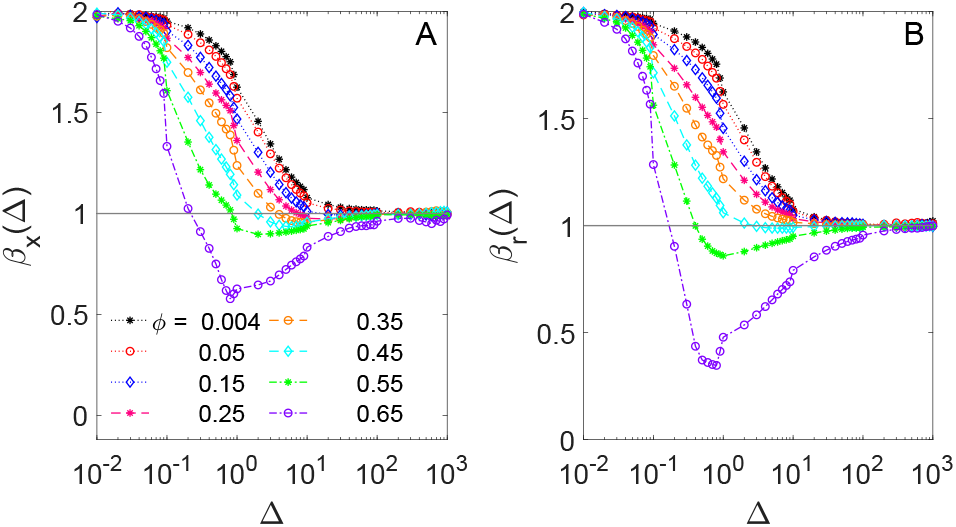
Time-local scaling exponents of (A) translational and (B) rotational TAMSDs (17) computed from the simulation data of Fig. 2 for varying crowding fractions *ϕ* of the dimers.

We emphasize that for star-shaped crowders a similar—but even more pronounced—non-monotonicity in the variation of these scaling exponents was detected at intermediate lag times, i.e., in the crossover region from the ballistic to the linear diffusion regime, see Fig. 6B in Refs.^93,94^.^║^ We find an equivalence of the ensemble- and time-based averaging for translational and rotational motion of the dimers. Namely, both in the short- and long-time regimes, we observe 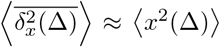 and 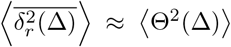 indicative of ergodic behavior in this Boltzmann-Khinchin sense^24,25^, see Fig. S4. This observation agrees with the results for the diffusion in crowded solutions of star-shaped crowders examined in Refs.^93,94^ (see also Sec. IVB below). We emphasize also an extremely small dispersion of individual TAMSDs around their mean, both for conditions of weak crowding (expected^156^ for massive-BM motion^15,16^) and heavy crowding, see the results in Fig. S4. This fact thus confirms the ergodicity of self-diffusion of dumbbells in crowded dispersions also in terms of the reproducibility of the TAMSD realizations^24,25^, for both translation and rotational motion.^**^

### B. Time-local scaling exponents

To examine the *ϕ*-dependence of the TAMSD in more detail, the time-local scaling exponents *β_x_* and *β_r_* are evaluated as^158^

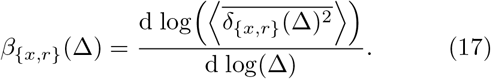

As follows from Fig. 3, crowding of the dimers has a profound impact on the behaviour of the scaling exponents at intermediate lag times, Δ ≈ 0.1… 10 × *δτ*. Namely, at very short times both scaling exponents *β_x_* and *β_r_* equal two for all crowding fractions of the dimers (the ballistic regime, see Eq. (6)), while for long lag times they both expectedly approach unity. In a dilute system, the crossover from the ballistic to the linear growth of the TAMSD occurs similarly to that for the MSD of a massive BM particle^15^, see also Eq. (16). Namely, the scaling exponent decreases monotonically from *β*_{*x,r*}_ = 2 at short to *β*_{*x,r*}_ = 1 at long times, see Fig. S2.

The translational and rotational motion of the dimers are correlated in terms of variations of their scaling exponents. We observe distinct *subdiffusive β_x_* and *β_r_* for the most crowded conditions, with the subdiffusion being more pronounced for rotational than for translational motion, compare the values of the dips at *ϕ* = 0.65 in Figs. 3A and 3B. These reduced values of *β_r_* at intermediate lag times and at high *ϕ* values are physically connected with a stronger impediment of rotational motion of a dimer by its neighbors, as compared to restrictions on possible translational motion of the same dumbbell.^††^

We refer the reader to Figs. 21 and 22 in Ref.^106^ for the analysis of 〈*x*^2^(*t*〉 and 〈Θ^2^(*t*)〉 in highly crowded 2D solutions of *active* dumbbells. This system also reveals nonmonotonic variations of the scaling exponents in the transitional regime after the initial ballistic growth, with more pronounced subdiffusion for rotations for *ϕ* ≳ 0.6.^‡‡^

### C. Diffusion coefficients

In the long-time limit, the exponents *β_x_* and *β_r_* converge to unity for all *ϕ* values, Fig. 3. Therefore, the fldependence of the mean TAMSDs 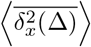 and 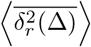 at long times stems solely from the *ϕ*-dependence of the respective translational *D_x_* and rotational *D_r_* diffusivities, evaluated in this regime of long-time BM as

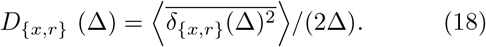

The fits of the TAMSD(Δ) are executed at lag times in the region Δ = 10^2^ … 10^4^ × *δτ*. The normalized diffusivities *D_x_* and *D_r_* as functions of varying crowding fractions *ϕ* are found to display similar behaviors. At small *ϕ*, both translational and rotational diffusivities decay exponentially,

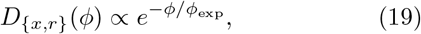

as evidenced in Fig. 4B.

**FIG. 4:**
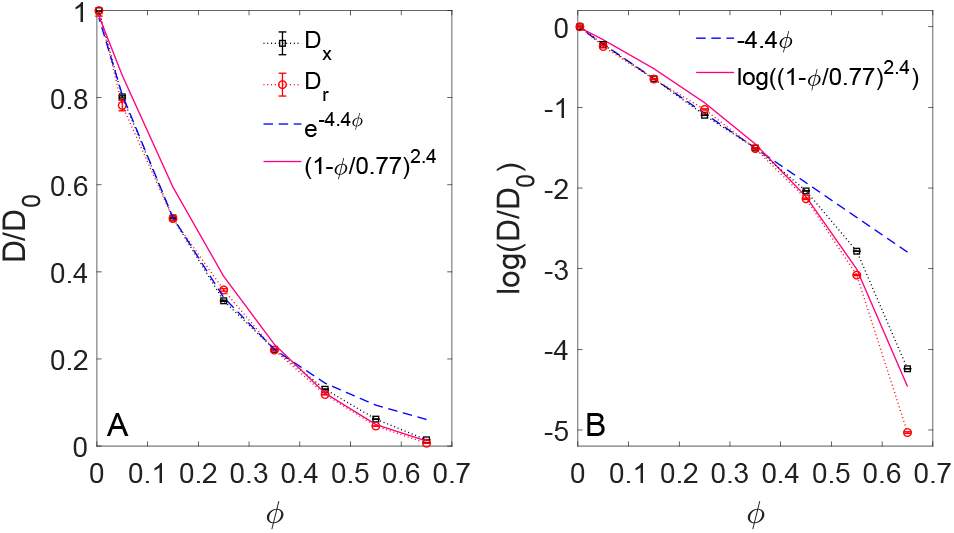
(A) Long-time translational and rotational diffusivities as functions of the crowding fraction *ϕ*, shown together with a fit to an exponential decay (19) for small **ϕ** and a quadratic fit (20) for large *ϕ* (see the legend). The values are normalized with respect to *D*_0_ = *D*(*ϕ* = 0.004) corresponding to a single dimer in the simulation box, see Tab. I. (B) Log_*e*_-scale representation of the simulation data from panel (A), with the same asymptotes shown.

The law (19) at small crowding fractions is also consistent with the linear-in-*ϕ* decrease of the diffusivity predicted theoretically and observed experimentally for concentrated suspensions of spheres in 3D, *D*(*ϕ*) ≈ 1 – *ϕ*/*ϕ*_exp_, see, e.g., Refs.^159,160^ and also Ref.^103,161^. A (slightly compressed) exponential decay of the tracer diffusivity in various hydrogel-like polymeric networks due to effects of steric obstruction was found and rationalized recently in Ref.^162^.

For large *ϕ*, we find a power-law dependence on the crowding fraction, see Fig. 4B. Similar dependencies were detected, e.g., for a system of polydisperse hard disks^77,163^ near the glass-transition point at nearly critical fractions of the crowding particles *ϕ*\*, namely

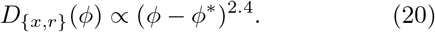

Similar values of the “critical exponent” were proposed for tracer self-diffusion in 3D hard-sphere suspensions, where *D*(*ϕ*) ∝ (*ϕ* – *ϕ*\*)^2^, see Refs.^164,165^. The dependence (20) is also in agreement with the results of mode-coupling theory^166,167^. We find that the magnitudes of the translational and rotational diffusivities are affected by varying *ϕ* values to a *similar* extent, in part because motions of disks imply also changes in orientation of the dimer (and, thus, changes in the angle Θ(*t*)). Of course, the relatively small scaling window available only allows for a relatively qualitative analysis.

This fact contrasts to some extent the properties of diffusion in the crowded solutions of three-arm stars studied in Refs.^93,94^, where the *D*(*ϕ*) ∝ (*ϕ* – *ϕ*\*)^2…2.4^ dependence was also detected. Namely, the translational self-diffusion of stars was found reduced more pronouncedly in crowded conditions, as compared to their rotational diffusion, see Fig. 7 of Ref.^93^. This is a physically expected trend because of a more isotropic overall appearance of the star-shaped crowders, as compared to the elongated, ellipsoidal-like shape of the dumbbells here. The translational motion for the stars and rotational motion of the dumbbells are, therefore, expected to be diminished stronger at a fixed high fraction of crowding.

In Fig. 4 we plot the dependencies (19) and (20) as the best fits in their respective ranges of fractions *ϕ*, realized for the parameters 1/*ϕ*_exp_ ≈ 4.4 and *ϕ*\* ≈ 0.77 for our computer simulation data (see the legend of Fig. 4). The crossover between these two apparent regimes—an exponential decay (19) in absence of subdiffusion (viscous) and a roughly quadratic dependence (20) for subdiffusion (viscoelastic)—takes place at *ϕ* ≈ 0.35… 0.45. Therefore, it occurs in the same range where the subdiffusive behavior for the TAMSD starts to emerge, see Fig. 3. We emphasize that—in contrast to the stronger subdiffusion for rotational motion of the dimers under conditions of moderate-to-severe crowding (see Fig. 3)— the translational and rotational diffusion coefficients are affected by self-crowding to a nearly the *same degree*, see Fig. 4.

We refer the reader to the studies of concentrated dispersions of (identical) spheres and of colloidal suspensions in 3D^74,159,160,164,165,168–177^ examining the effects of hydrodynamic interactions. In a number of these studies, the dependence of the long-time self-diffusion coefficient of the spheres [with hydrodynamics] was proposed to have a quadratic form,

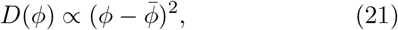

comparing well with experiments^165,172^, see Eq. (20). Hydrodynamic effects were considered in detail also in Refs.^65,176^; we also refer to the recent comparative analysis^177^ and to Refs.^161,175,178–180^ for the examination of translational and rotational diffusion in dense suspensions of charged colloids, see also Ref.^176^. A detailed experimental study of self-diffusion in concentrated solutions of *bovine serum albumin* proteins—including an in-depth discussion of the *D*(*ϕ*)-dependence in the presence of hydrodynamic and electrostatic interactions—can be found in Ref.^110^.

### D. Dimer-displacements PDF

The PDFs of translational *p*(*δ_x_*) and rotational *p*(*δ_r_*) displacements of dumbbells, computed in the range Δ = 0.1… 80 × *δτ*, are plotted in Fig. 5 for the most crowded system at *ϕ* = 0.65. At short lag times, for Δ = 0.1 × *δτ*, the PDFs for both types of motion expectedly have Gaussian-like profiles. At longer times, the translational PDF remains nearly Gaussian, while the rotational PDF becomes nearly Laplacian (e.g., exponential, as quantified in Sec. IIIE below). This feature is visible from the nearly straight tails of the PDF in log-linear scale, as presented in Fig. 5B. This fact indicates that the rotations of dumbbells on large azimuthal angles are much more frequent than the expectation from a Gaussian form of the rotational PDF. We stress here that the statistical observables examined for the crowded solutions of dumbbells in Secs. III D, III E, III F, and III G are new compared to the analysis of self-diffusion of star-shaped crowders performed in Refs.^93,94^.

**FIG. 5:**
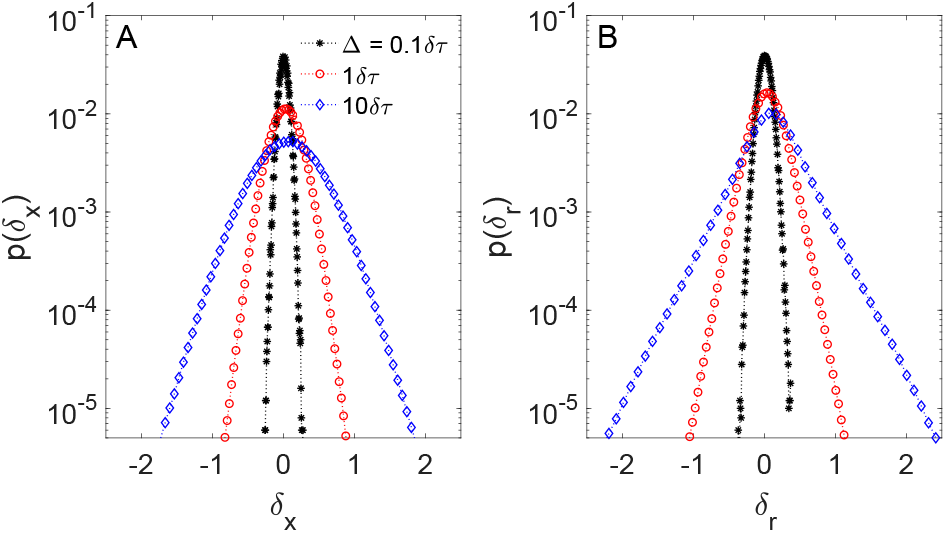
(A) Translational and (B) rotational displacement-PDFs for the highest crowding fraction *ϕ* = 0.65, computed for several values of the lag time Δ (see the legend) and for other parameters (such as *m, γ, τ*_0_, *ϵ*, and *T*) as in Fig. 2.

**FIG. 6:**
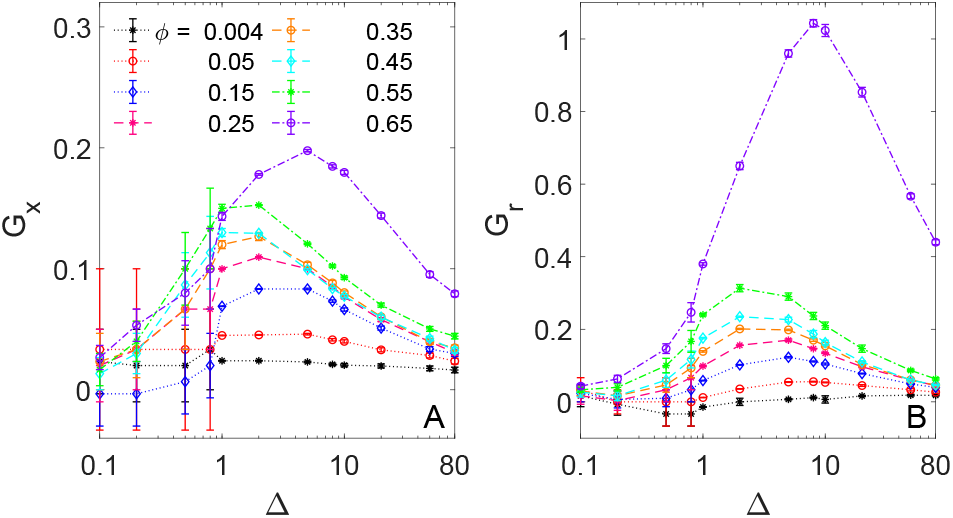
Non-Gaussianity parameter (22) for (A) translational and (B) rotational motion of the dumbbells plotted against the lag time Δ for a set of varying *ϕ* (see the legend). The lines connecting the points guide the eye. Note that panels (A) and (B) have different vertical scales.

**FIG. 7:**
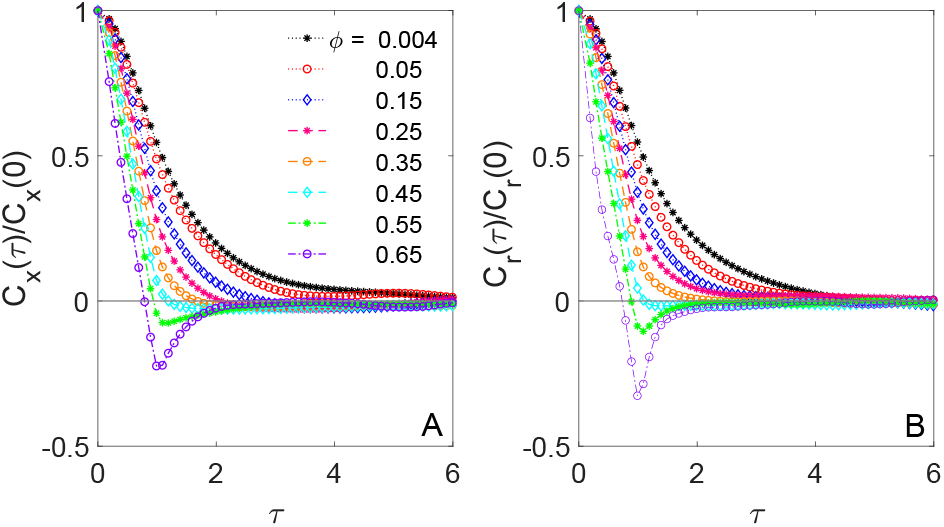
Normalized (A) translational and (B) rotational displacement ACFs as a function of the correlation time *τ* computed for varying crowding fractions *ϕ* (see the legend). The presented ACFs are normalized to the respective initial values at *τ* = 0. Parameters: the lag time in Eq. (24) is Δ = 1 × *δτ*.

### E. Non-Gaussianity parameter

The degree of non-Gaussianity of the PDF shapes of both translational and rotational displacements of the dumbbells is quantified in terms of the non-Gaussianity parameter *G*^27^. It is related to the kurtosis (denoted as Kurt below) of the PDF distributions of the displacements of the dimers as

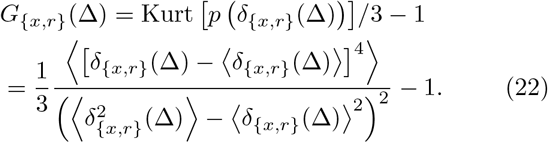

Here, the *n*th moment of the PDF of the displacements *p*(*δ*_{*x,r*}_)—denoted as 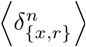 below—is defined for natural numbers *n* ≥ 1 as

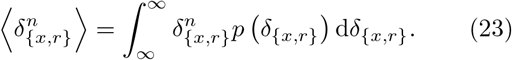

Thus, in expressions (22) and (23) the ensemble-averaged moments are used, as compared to the TAMSDs given by Eqs. (14) and (15).

In Fig. 6 we present the lag-time dependencies of the non-Gaussianity parameters (22) for varying crowding fractions *ϕ*. Corroborating the Gaussian PDFs at short times (see Fig. 5) the values of *G*_{*x,r*}_ are comparatively small in this regime. The values of *G_x_* and *G_r_* increase and reach a maximum at intermediate lag times. At long lag times, extending to the regime of BM for the respective MSD and TAMSD, the values of *G_x_* and *G_r_* tend to relatively small values again. These nearly Gaussian PDFs for the displacement distributions obtained from our simulations at very long times—in the BM-regime of the MSD at Δ = 80 × *δτ*, as shown in Fig. 6—are also expected theoretically from the central limit theorem. Note here that the longest time Δ = 10 × *δτ* for the computed rotational displacement-PDF in Fig. 5B is still considerably shorter than the longest time Δ = 80 × *δτ* in the plot of Fig. 6B for the respective computed *G_r_*.

As expected, the most pronounced deviations from Gaussianity are detected for the displacement distributions *p*(*δ*_{*x,r*}_) at the maximal crowding fraction, *ϕ* = 0.65. The maximum of the translational non-Gaussianity parameter is *G_x_* ≈ 0. 2, while the maximum of the rotational non-Gaussianity parameter is significantly larger, *G_r_* ≈ 1. The latter fact indicates again a nearly Laplacian shape of the rotational-displacements PDF illustrated in Fig. 5B.^§§^ We also demonstrate in Fig. S5 that the second moments for the PDFs *p*(*δ*_{*x,r*}_)(Δ))— when computed from the simulation data shown in Fig. 5—reveal similar features in their lag-time dependencies in the range Δ = 0.1… 80 *δτ* for varying *ϕ* fractions, as those of the respective TAMSDs 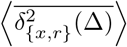 shown in Fig. 2A,B.

### F. ACF of displacements and rotations

We now evaluate the velocity-ACF *C_x_* for the translational displacements *δ_x_*, rotational displacements *δ_r_* and the displacements *δ_d_* of the relative coordinate in the dimers. With the time difference *τ* between the two displacements—the correlation time—the ACF is obtained as the time average,

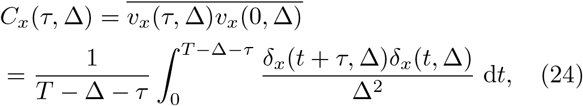

where the factor 1/Δ^2^ stems from the definition of the “velocity”, *υ_x_*(*t*) = [*x*(Δ + *t*) – *x*(*t*)]/Δ ≡ *δ_x_*(*t*, Δ)/Δ. The dependencies of the time-averaged ACFs for rotational motion and relative-displacement motion of the disks in a dimer, *C_r_* and *C_d_*, respectively, are calculated analogously to expression (24). As the displacements in the integrand of Eq. (24) are shifted by *τ*, the time averaging is executed along the recorded trajectories up to the trace length (*T* – Δ – *τ*).

In Fig. 7, we present the translational and rotational displacement-ACFs plotted against the correlation time *τ* for varying *ϕ*. For weak-crowding conditions, the decay of the displacement-ACFs *C_x_* and *C_r_* is monotonic, while for very crowded systems the ACFs display a nonmonotonic behavior. Specifically, initially at *τ* ≤ Δ the normalized ACFs start decreasing from unity, reaching a minimum at *τ* = Δ. At later stages, for longer correlation times *τ*, the “recovery” of the ACFs to zero from the region of negative values is detected in the simulation data, see panels (A) and (B) of Fig. 7 for translation and rotational motions, respectively. The displacement-ACFs for larger Δ values presented in Fig. S7 clearly reveal a trend of progressively *shallower* minima at later Δ.^¶¶^

These anticorrelations at *τ* = Δ in the highly crowded solutions physically indicate a likely reversal of motion of the dumbbells in the consecutive time step, often referred to as antipersistence. The form of the displacements-ACFs *C_x_* and *C_r_* themselves are reminiscent of those for subdiffusive processes of fractional BM and of motion governed by the fractional Langevin equation^24–26,31,32^.

These two models of non-Brownian diffusion are often used for the mathematical description of subdiffusion in crowded and viscoelastic media^26,184^.^***^

### G. Relative-coordinate ACF

The normalized ACF *C_d_*(*τ*)/*C_d_*(0) for displacements of the relative coordinate—defined in the time-averaged sense as in Eq. (24)—is plotted versus *τ* for *ϕ* = 0. 004 (a single dimer in the simulation box) in Fig. 8. The oscillations of *C_d_*(*τ*) indicate vibrational “breathing modes” of the dimers and the exponentially decaying amplitude of these oscillations is due to the damping effects of the environment. As effective damping increases with *ϕ*, the oscillations become suppressed and their amplitudes decay faster with time as *ϕ* grows, compare the results of Fig. 8 for a single dimer to those of Figs. 9 and 10 obtained for crowding fractions *ϕ* = {0.05, 0.15,0.25,0.35}.

**FIG. 8:**
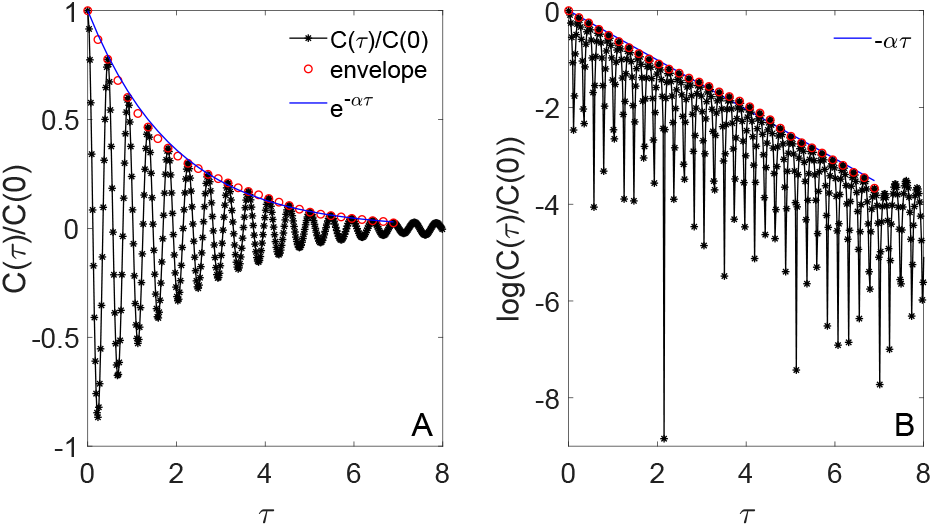
(A) Normalized displacement ACF *C_d_* of the relative coordinate of a single dimer (*ϕ* = 0.004) as a function of the correlation time *τ*. This crowding fraction corresponds to a single dimer in the simulation box, see Tab. I. The decay of the ACF amplitude is fitted by the envelope function 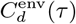 given by (25). (B) Data from panel (A) in log_*e*_-scale (shown together with a linear asymptote for the same exponential decay).

**FIG. 9:**
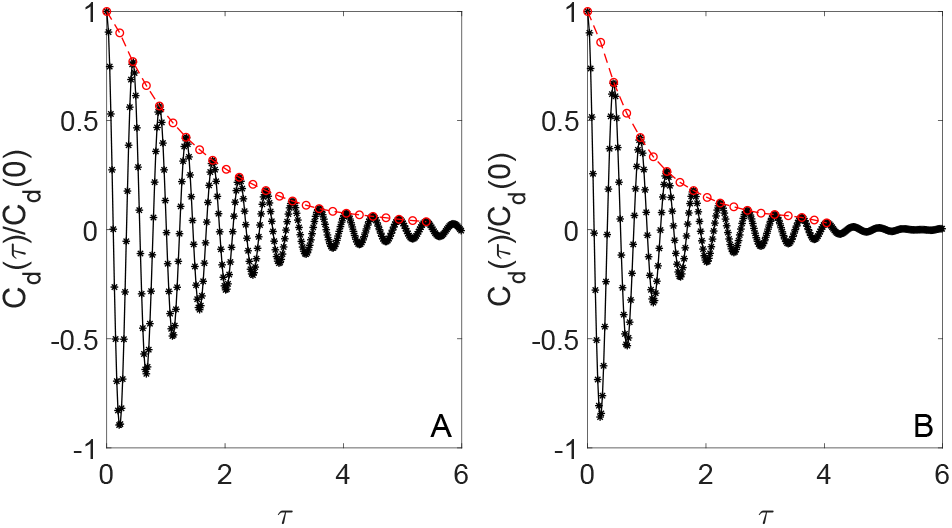
Displacement ACF *C_d_* for (A) *ϕ* = 0.05 and (B) *ϕ* = 0.15. The red circles are given by Eq. (25).

**FIG. 10:**
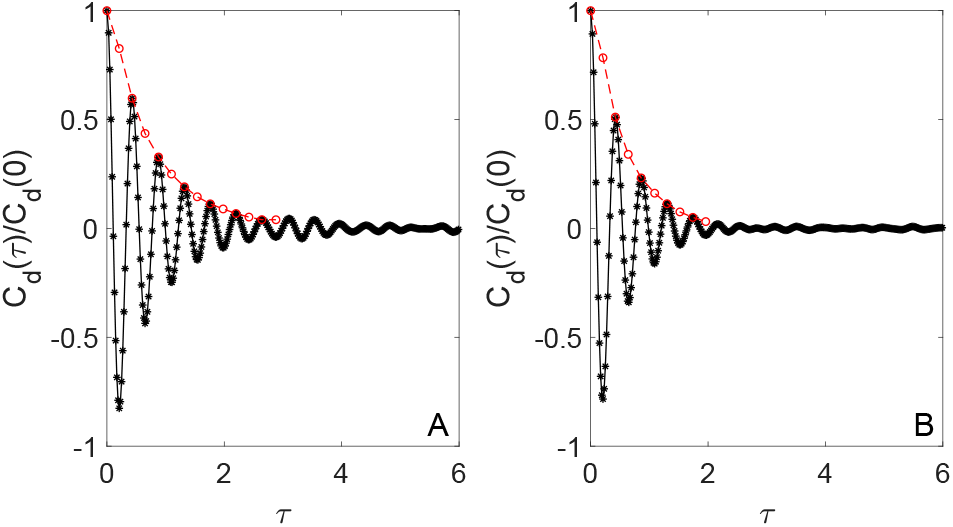
The same as in Fig. 9, but computed for (A) *ϕ* = 0.25 and (B) *ϕ* = 0.35. The red circles are given by Eq. (25).

We compare these oscillations of the dumbbells to those of a noise-driven damped harmonic oscillator (see §25 of Ref.^185^). The period of *C_d_*(*τ*) oscillations, denoted by T, is found to be only moderately dependent on *ϕ*, see Fig. S8A. For the frequency of dimer oscillations, *ω* = 2*π*/T, for weak crowding [and no effects of the neighbors] we find an excellent agreement with the eigenfrequency of the damped harmonic oscillator^16,185^ *ω*_0_ = 2*k/m* [for free oscillations, with the reduced mass *m*/2].

With increasing *ϕ* the simulation data predict *increasing* frequency of *C_d_*(*τ*) oscillations. Physically, this trend is expected because—for a given energy “ influx” into the system due to fluctuations of the thermal bath—the reduced space available for expansion of the dumbbells in progressively more crowded solutions gives rise to oscillations at higher frequencies. The frequency of the damped oscillator, however, has the *opposite* trend with increasing crowding fraction [if we assume that *α*(*ϕ*) ∝ *γ*(*ϕ*), as in Fig. S8B]. In both cases, however, rather insignificant *ω*(*ϕ*) variations are found, as quantified in Fig. S8.

The *ϕ*-dependence of the decay rate *α*(*ϕ*) of the envelope function 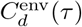 of the oscillating ACF-function *C_d_*(*τ*) is determined from the exponential fit

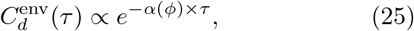

as shown in Fig. 11. To relate α(*ϕ*) to the exponential dependence of *D*(*ϕ*) in Fig. 4 and Eq. (19), the normalized *inverse* decay constant is shown in Fig. 11B. We find that, similarly to *D*(*ϕ*), the reciprocal constant 1/*α*(*ϕ*) decays nearly exponentially with *ϕ* and only the most crowded solutions reveal deviations from this behavior. The inverse representation was chosen in Fig. 11A in order to compare the resulting dependence of *α*_0_/*α*(*ϕ*) to a decreasing *D*(*ϕ*) in Fig. 4A [because *α*(*ϕ*) ∝ *γ*(*ϕ*) and *γ*(*ϕ*) ∝ 1/*D*(*ϕ*) in virtue of relation (3)]. The decaying oscillations of the displacement-ACF *C_d_*(*τ*) of the dumbbells are typical for a damped harmonic oscillator^16,185^. This similarity is expected because Eq. (10) reduces to the Langevin equation of a randomly driven harmonic oscillator when a single dimer is considered. In the dilute regime the Lennard-Jones interactions can in the first order be neglected because the distance between the two monomers is ~ *r*_0_. In addition, as proposed in Ref.^16^, it gives a physical meaning to the decay rate *α* ~ 1/*τ*_0_. Therefore, the vibrations of the dimers for *weakly crowded* systems can be rationalized by those of randomly driven harmonic oscillators with the crowding-fraction dependent damping constant, *γ*_eff_(*ϕ*).

**FIG. 11:**
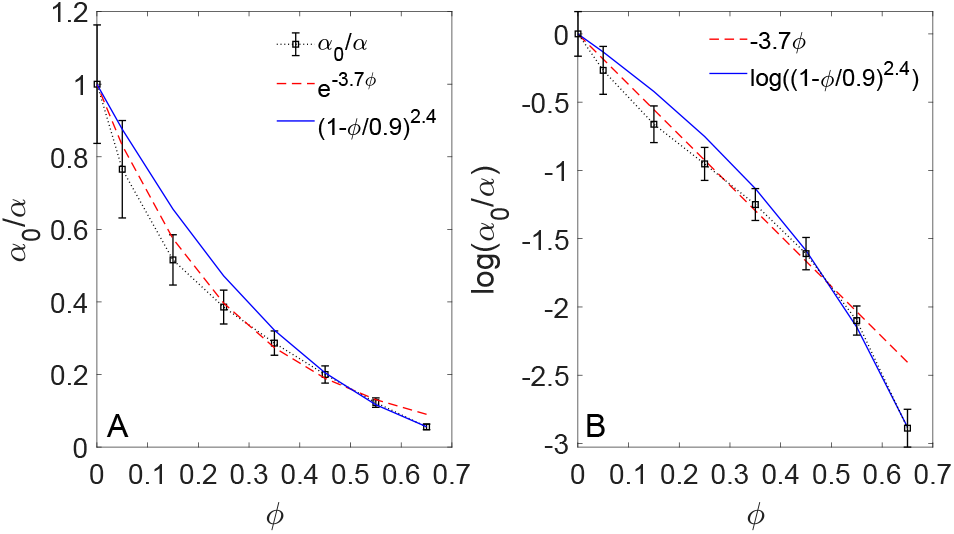
Normalized inverse decay constant of 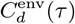 given by *α*_0_/*α* from Eq. (25) shown as function of crowding fraction *ϕ* in linear (A) and log_*e*_ (B) scale. The fit functions used are listed in the legends. Here, *α*_0_ = *α*(*ϕ* = 0.004) and the error bars are indicated.

We note that some deviations from the behavior ( 25) observed at large *ϕ* in Fig. 11 indicate a potential change to another diffusion regime. Due to worsening accuracy of the simulation data for *α* in this regime, however, the crossover is not clear. This high-density regime may be modeled by a fractional Langevin equation with powerlaw memory kernel in the presence of an external harmonic potential^32,35^. To be confident about these observations significantly more elaborate simulations are needed.

## IV. DISCUSSION AND CONCLUSIONS

### A. Summary of the main results

Based on the results of extensive computer simulations we studied the stochastic dynamics of a crowded system of dimers in two distinct regimes of crowding fractions *ϕ*. In the region of small *ϕ*, at *ϕ* ≲ 0.35, we observed standard BM of the dimers supported by the behavior of the scaling exponent in Fig. 3 and by variation of the displacement-ACF of the COM of the dumbbells shown in Fig. 7. Note that the deviations of the ACF from zero at larger *τ* values are due to progressively worsening statistics. In this weak-crowding regime, the effects of crowding can be modeled via an effective constant *γ*_eff_, compare Figs. 3 and S2. Using Eq. (3), the variation of *γ*_eff_(*ϕ*) obtained from Fig. 4 is, therefore, also nearly exponential with *ϕ*, see expression (19).

At high-crowding conditions, with *ϕ* ≳ 0.35, we observed subdiffusive scaling exponents (with particularly severe subdiffusion for rotational motion) and the displacement-ACFs displaying anticorrelations indicative of, e.g., viscoelastic-type subdiffusion^24^. We found that translational and rotational motion of the dimers are coupled in the region of intermediate lag times where crowding-induced subdiffusion is detected, see Fig. 3 and also Refs.^91,102^. In this regime, with increasing crowding fractions *ϕ* the diffusivity decreases according to a powerlaw form, Eq. (20).

A model that combines subdiffusion and anticorrelations is subdiffusive fractional Brownian motion^186^ as well as the closely related model of motion governed by a fractional Langevin equation^187^. These two mathematical models of anomalous but ergodic diffusion are often associated with the physical motion of [endogenous and exogenous] tracers in viscoelastic environments^24,52,128^, such as those of the cell cytoplasm and of artificially crowded fluids *in vitro*. Thus, we can interpret the two distinct regimes regarding the observed effects of crowding on the diffusion of the dimers as those of a viscous liquid at low *ϕ* and viscoelastic liquid at high *ϕ* fractions. A concentration-dependent transition between viscous and viscoelastic diffusion was also proposed in Ref. ^188^.

### B. Physical rationales and further discussion

We found two distinct diffusion regimes in our crowded system of passively diffusing dumbbells. In dilute systems, we obtained a behavior consistent with standard BM featuring Gaussian displacement-PDFs and nearly exponentially decaying (Laplacian) rotational ACFs. The effect of *ϕ* on the diffusion could be modeled here as effective damping, obtained from the exponential decrease of the diffusivity with *ϕ*, see Eq. (19). Similarly, the damping of internal vibrations of the dimers was shown to depend exponentially on *ϕ*. It was found to give rise to a quicker decay of oscillations in the respective internal-ACF describing relative separations of monomers in a given dumbbell.

For highly crowded systems, we instead observed transient subdiffusion of the dumbbells and the non-Gaussian PDFs of their anticorrelated displacements. In this regime, the average diffusivity of the dumbbells was shown to decrease as a power law with increasing crowding fraction *ϕ*, Eq. (20).

We tentatively attribute these two different diffusion regimes found in simulations to a crowding-induced crossover from a purely viscous to a viscoelastic behavior of the diffusion environment. Physically, this viscoelasticity at high *ϕ* can stem from shape-responsiveness and “concerted” motions of neighboring dumbbells. Namely, noise-driven “jiggling” of dimers leads to highly frequent collisions and interactions between them. A finite adaptability of elastically responsive dumbbells can thus give rise to *correlated* shape deformations of the neighbors at high *ϕ* values. A displacement of a dimer COM position from its equilibrium value gets reversed by the elastic environment of the neighbors leading to anticorrelated motions of a given dumbbell, see Fig. 7. To verify this hypothesis, further studies should be done to include a *ϕ*-dependence of inter-monomer forces and for different number of neighbors.

Recent Bayesian model-assessment analysis^189–191^ as well as modern machine-learning approaches^192–198^ could then determine relative probabilities of possible diffusion models involved. In the end, certain predictions regarding the diffusion characteristics based on physical properties of the environment of crowders might be possible.

We also observed this transition for the diffusion of *single non-connected* monomers characterized by a crossover of the diffusivity dependencies, see Fig. S6. Here, the transition occurs at somewhat higher *ϕ* values because solely the Lennard-Jones potential is the source of elasticity in this modified system. This suggests that more complex crowders—such as a linear chain of inter-connected monomers^97^—should exhibit such a transition from viscous to viscoelastic behavior at smaller *ϕ* fractions, as compared to those for the dumbbells (work in progress).

To discuss the effects of the dumbbell shape on the diffusive characteristics, we highlight now the key differences of the current results to those for the diffusion of star-shaped crowders reported earlier in Ref.^93^. For the star-shaped crowders, transient subdiffusion was found to occur for *all* crowding fractions (up to *ϕ* = 0.55), while only the most crowded dimer-based systems were shown to exhibit such a behavior, see Figs. 2 and 3. Similarly, the power-law dependence of the diffusivity on the crowding fraction, *D*(*ϕ*), was observed in Ref.^93^ for *all ϕ* values, while for the dimers we observed this dependence only in a high-*ϕ* regime. Moreover, the crowding fraction corresponding to the glass transition value *ϕ*\* for the star-shaped crowders^93^ was smaller than that for the dumbbell-shaped crowders examined in this study, 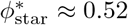 versus 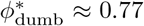, respectively. For starshaped crowders^93^, at “intermediate” time scales strong variations of both super- and subdiffusive behaviors were observed^94^, with a more anomalous TAMSD scaling exponent detected for rotational than for translational motion, similarly to the dimers. Moreover, the diffusion of star-shaped crowders was found ergodic for small *ϕ* values^93^.

These deviations can, in part, be due to different structures of the two types of crowders. The inner monomer of a star-like crowder^93,94^ is connected to three other monomers by springs and, thus, its environment is effectively “viscoelastic” independent of *ϕ* value. In contrast, a disk in a dumbbell is connected to only one other monomer, allowing for a more “viscous” medium at low *ϕ*. The difference of 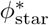 and 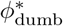 can be explained by a simpler structure of dimers facilitating their denser packing prior to reaching the glass-transition point.

Although an elastic dimer can be considered as a model of shape-asymmetric proteins^132,199^, 3D computer simulations with electrostatic interactions and hydrodynamic effects^63^—instead of 2D simulations without these two effects—are required to approach the problem of macromolecular crowding as realized in many biological cells. Crowding by shape-elongated molecules can have implications onto the properties of biomolecular reactions^66^ (including protein folding and protein-protein association) as well as of polymer and DNA looping. The diffusion in the crowded solutions of anisotropic proteins in lipid membranes^100,101^ and the elongated particles (such as short fragments of DNA and rod-like viruses) adsorbed on lipid membranes^200–205^ are also relevant areas and biophysical systems.

Finally, size- and shape-asymmetric particles (charged dumbbells^76^ and neutral molecules) can also build some constituents of ionic liquids^126,127,206,207^. Dumbbell-shaped molecules functionalized with ionic liquids were proposed, e.g., as “hybrid” electrolyte for lithium-metal batteries^208^. Soft dumbbell particles with opposite charges^209–212^ and their applications to dipolar liquids and gels can also be mentioned.

## Acknowledgments

A. G. C. gratefully acknowledges the Humboldt University of Berlin for hospitality and support. The authors thank D. Caetano, S. Kondrat, and R. G. Winkler for correspondence and scientific discussions/comments. R. M. acknowledges financial support by Deutsche Forschungsgemeinschaft (DFG Grant ME 1535/12-1). R. M. thanks the Foundation for Polish Science (Fundacja na rzecz Nauki Polskiej) for support within an Alexander von Humboldt Polish Honorary Research Scholarship.

## Abbreviations

BM: Brownian motion
COM: center of mass
PDF: probability-density function
MSD: mean-squared displacement
TAMSD: time-averaged MSD
ACF: autocorrelation function

## Appendix A: Simulation scheme

To simulate Eq. (10), we use the numerical Verlet velocity algorithm for the positions and velocities of the *i*th particle. We use it separately for the *x*- and *y*-component, but present below only the expressions for the *x*-component, namely

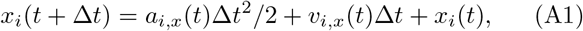

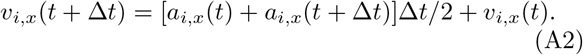

The integration time-step is Δ*t* = 0.005. For the acceleration of the *i*th disk, with *x*-component *a_i,x_*(*t*), we use the forces from the potentials (8) and (9) yielding

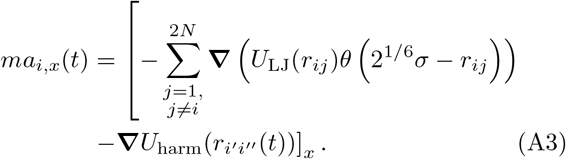

We describe friction via the *γ*-dependent coefficients *c*_{*x*1,*x*2}_(Δ*t*) and *c*_{*υ*1,*υ*2,*υ*3}_(Δ*t*) given by (see Ref.^150^)

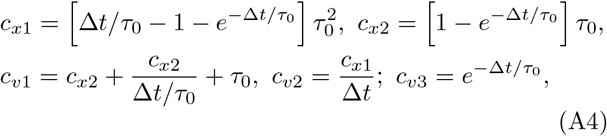

so that

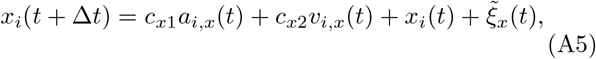

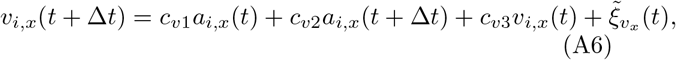

where the noise is modeled by random variables 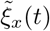 and 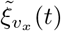, At *γ* → 0—that physically corresponds to the limit of undamped motion and of the Newtonian^13^ dynamics—expressions (A5) and (A6) turn into (A1) and (A2).

## Appendix B: Supplementary figures

**FIG. S1:**
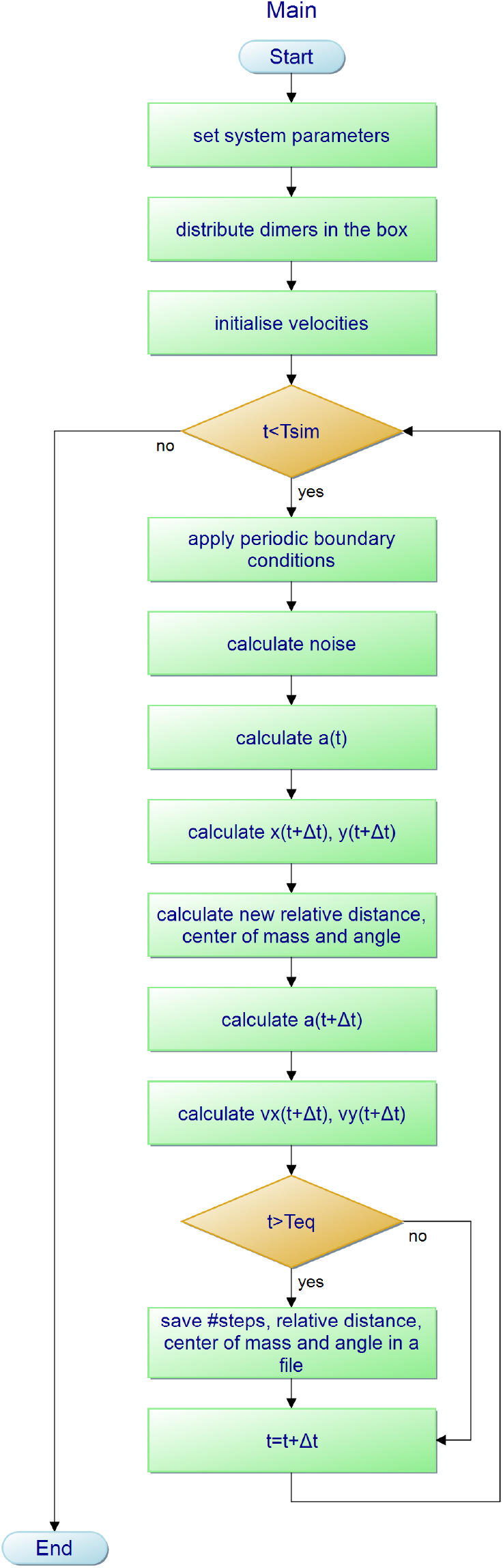
Flow-chart of computer simulations, with initialization of the system and integration loop.

**FIG. S2:**
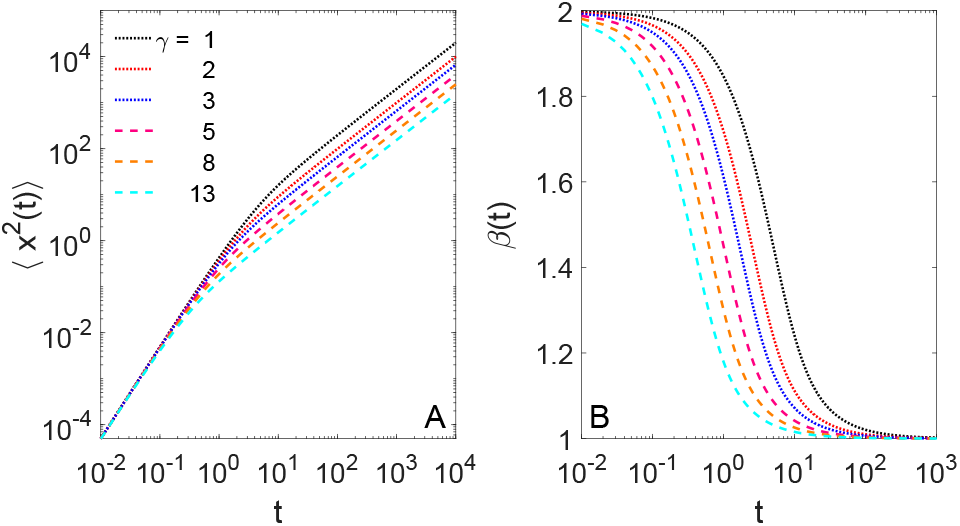
(A) Translational MSD for the COM motion of a single dimer governed by Eq. (4) (see also Eq. (17) in Ref.^152^) for varying friction coefficient *γ*, with 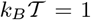 and *m* = 1. (B) MSD scaling exponent for the simulation data of panel (A), calculated via Eq. (17).

**FIG. S3:**
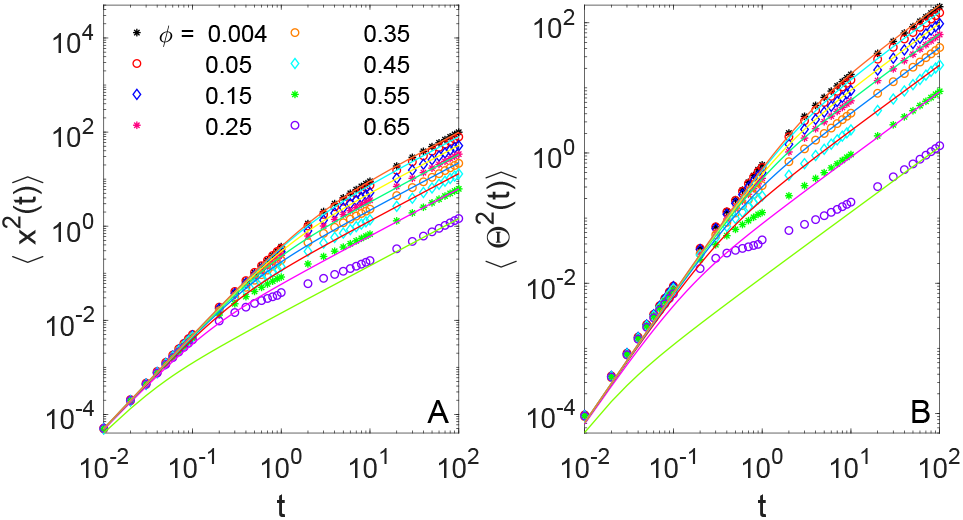
(A) Translational and (B) rotational MSD for diffusion of dimers at different crowding fractions *ϕ* (see the legend, symbols), plotted for *γ* = 1 for the TAMSD data of Fig. 2, with the analytical result (16) shown as the solid curves.

**FIG. S4:**
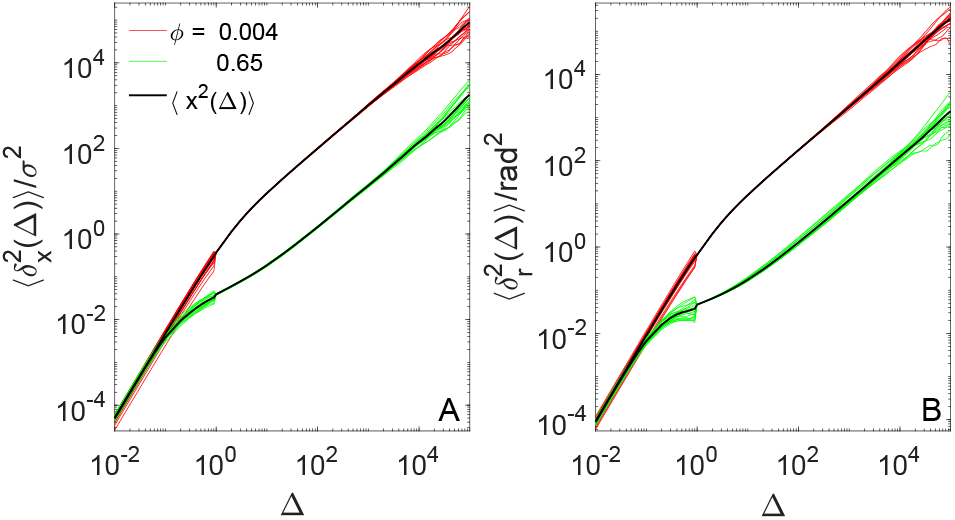
Direct comparison of the translational (A) and rotational (B) MSDs (the solid black curves) and the TAMSDs (the colored curves) for self-diffusion of dumbbells at two limiting crowding fractions, *ϕ*_min_ = 0.004 and *ϕ*_max_ = 0.65 (yielding fast and slow diffusion, respectively, see the legend). The results are plotted from the TAMSD data of Fig. 2 and the MSD data of Fig. S3. Note that, as the integration lag-time interval was splitted into two domains to facilitate the performance, a minor discontinuity at Δ = 1 × *δτ* is visible in the data.

**FIG. S5:**
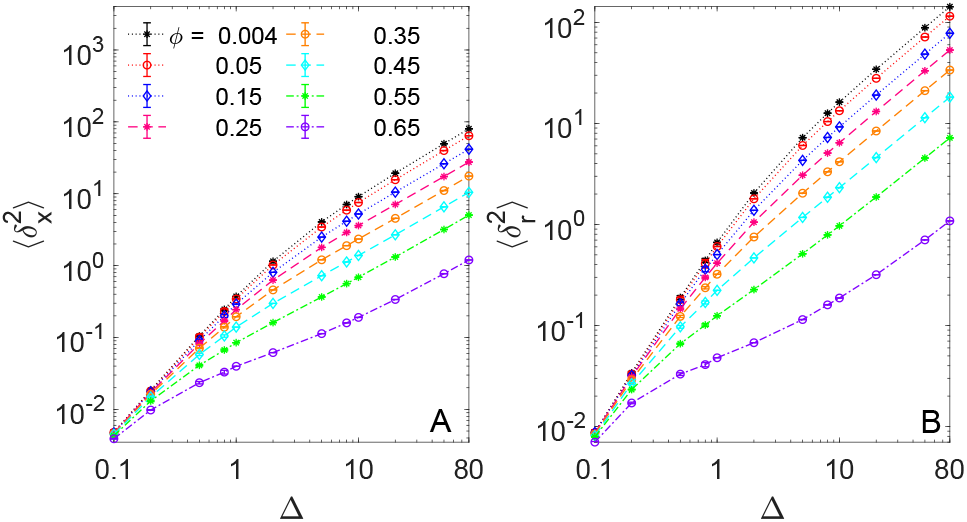
Second moment of the (A) translational and (B) rotational displacement-PDFs given by Eq. (23), presented as a function of lag time Δ for varying crowding fractions *ϕ*.

**FIG. S6:**
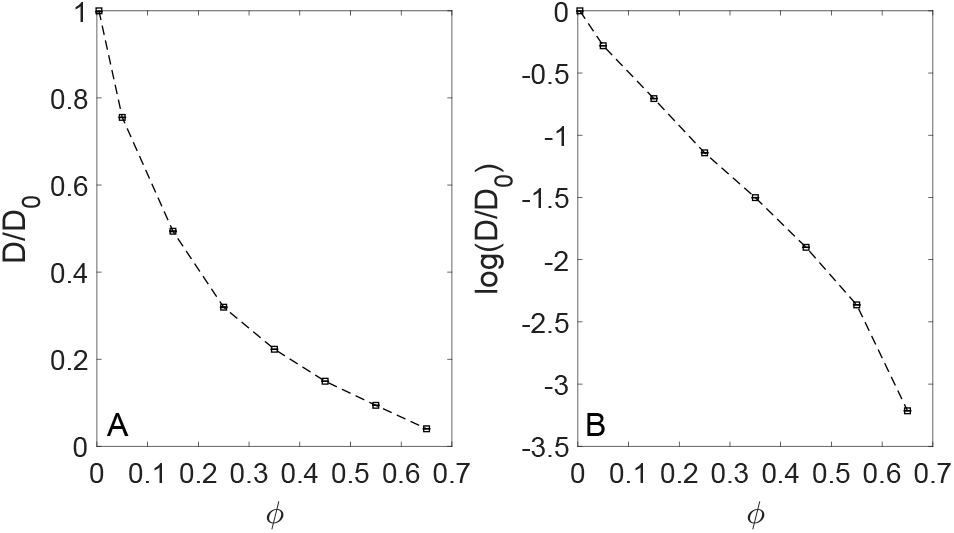
(A) Translational diffusivity as a function of *ϕ* obtained via expression (18) from the MSD of diffusion of single, non-connected monomers. (B) Linear-log_*e*_ plot of the data of panel (A). Parameters are the same as in Fig. 2.

**FIG. S7:**
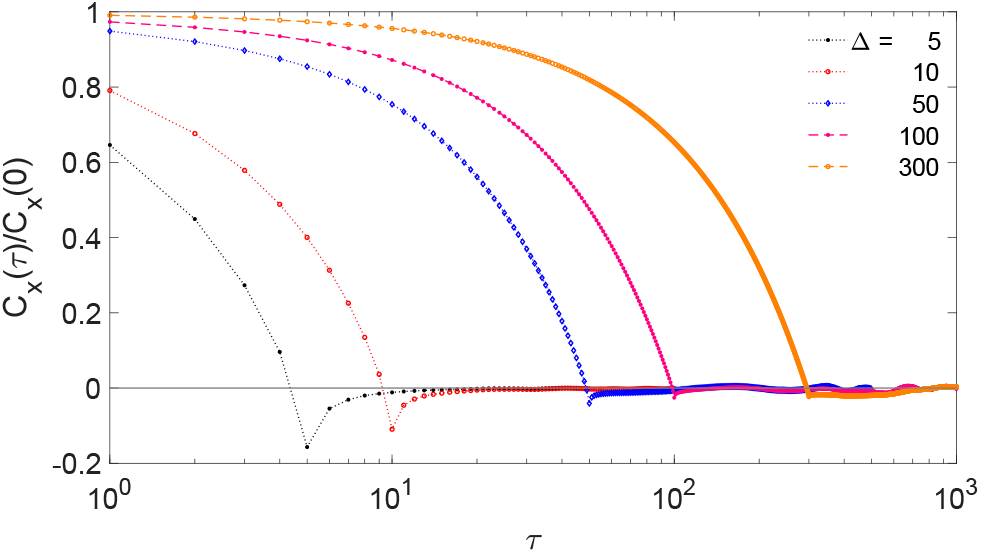
The same as in Fig. 7, evaluated for the same parameters except for larger values of the lag time, see the legend. All the curves approach unity at vanishing *τ* values (the region is not shown in the plot for presentation purposes).

**FIG. S8:**
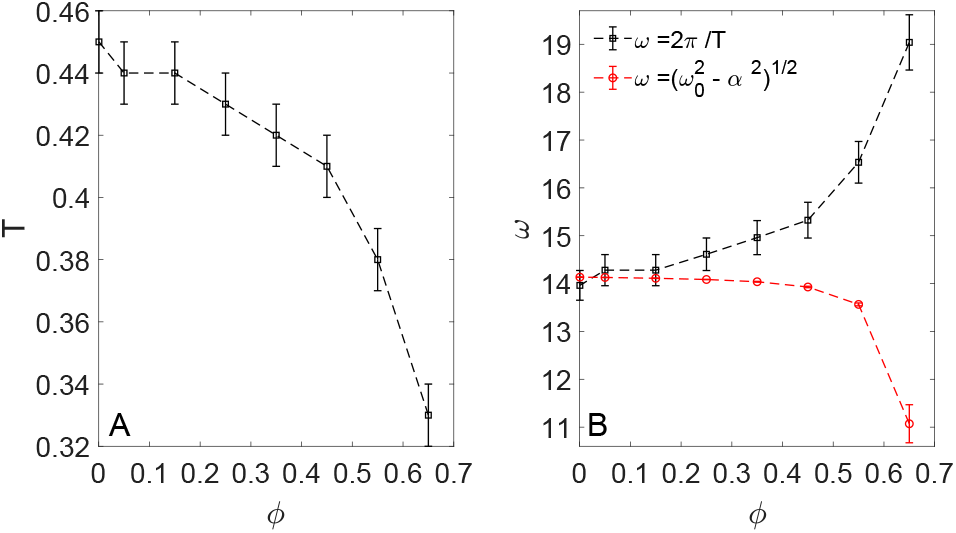
(A) Variation of the oscillation period T(*ϕ*) of the ACFs *C_d_*(*τ*)—as obtained from the simulation data of Figs. 8, 9, and 10—versus crowding fraction *ϕ*. (B) Comparison of the recalculated frequencies *ω*(*ϕ*) = 2*π*/T(*ϕ*) plotted versus *ϕ* to the frequency of the damped harmonic oscillator^16,185^. The latter is given by 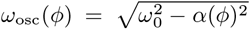, where the eigenfrequency is *ω*_0_ and we assumed *α*(*ϕ*) ≫ *γ*(*ϕ*) ≫ 1/*D*(*ϕ*).

## Footnotes

* As an example, superdiffusion detected in the cytoplasm of *Acanthamoebae* was attributed to persistent active motions caused by myosin-II motors and cell locomotion^51^. Conversely, subdiffusion is often associated with the passive motion of tracers in crowded, polymer containing, “restrictive” environments giving rise to antipersistent motion, such as the cytoplasm of biological cells^52^.

† Here, the *r*^−12^-term models a strong short-range repulsion [stemming from the quantum-mechanical exclusion in overlapping electron clouds], whereas the *r*^−6^-term mimics a weak long-range vander-Waals-like attraction, as in the models of diatomic molecules and chemical bonds^133–136^.

‡ Note that the maximal packing fraction of identical disks in 2D is^141–143^ 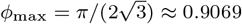 (the hexagonal packing), while in 3D the maximal packing density of identical spheres is (see, e.g., Ref.^144^) 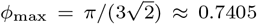 (the so-called “cannon-ball” packing).

§ Note that, as in Ref.^93^, the time *T*_cr_ grows with *ϕ* because *D*(*ϕ*) is a decreasing function of the crowding fraction *ϕ* (as we demonstrate below). Therefore, the most crowded system is taken below as a “reference” for the characteristic diffusion time, namely *T*_cr_ = *T*_cr_(*ϕ*_max_ = 0.65).

¶ Note that for the rotational diffusion the cumulative angle is computed in the simulations and, thus, the angle difference is *not* bounded by 2*π*. Therefore, for *n* full rotations of a dumbbell in units of panel (B) of Fig. 2 the azimuthal MSD amounts to (2*πn*)^2^.

║ Note that a nonmonotonic variation of the TAMSD scaling exponent with lag time was also detected, e.g., for anomalous diffusion of proteins and lipids on/in lipid membranes^130,131^ and in the model of tracer diffusion in driven lattice Lorentz gases of immobile obstacles^154,155^.

** We do not quantify here the spread of individual TAMSDs and the respective dispersion determining the value of the ergodicity breaking parameter^24,25,93^. We assess ergodicity^157^ of the system *solely* from the equivalence of the MSD and the mean TAMSD at short lag times.

†† Note that variation of the moment of inertia of the dumbbell— e.g., via altering its mass distribution and size—one can tune the rotational relaxation time *To* θ. Coincidentally, for the currently chosen model parameters the regions of the TAMSD-based subdiffusion for translational and rotational motions are observed at nearly the *same* lag times, see panels (A) and (B) of Fig. 3. Simulating dimers with the same geometric shape and total mass but with significantly different moment of inertia *I*—positioning, e.g., artificially the whole mass in the central point of the dimer and thus making *I* = 0—could also be performed. This would answer the question regarding coupling and inter-relation of the diffusion properties (the region of subdiffusion, values of the scaling exponents, etc.) for translational and rotational motion of crowded solutions stemming from the inertia of the dumbbells versus from correlations in their motion.

‡‡ Note that the diffusion of 2D dimers has recently also been considered experimentally and theoretically in Ref.^102^, with the focus on translational-rotational coupling. Specifically, at low *ϕ* ≈0.14 realized in experiments^102^ the diffusion of nearly independent dimers was assumed. In contrast, our dumbbells are often at high *ϕ*, when the assumption of “independence” is violated. In addition to obtaining the low-order moments, a generalized scattering function was derived^102^. It was found that translational-rotational coupling is caused by *anisotropic* diffusion of the dimers parallel and perpendicular to their axis, also detected in granular gases^123^. While we observe a similar coupling of motion, see Fig. 3, no longitudinal and transversal diffusivities of the dumbbells were recorded in current computer simulations.

§§ We remind the reader here that Kurt=3 for a Gaussian PDF in 1D, while Kurt=6 for a Laplacian distribution. Therefore, Eq. (22) gives for these two cases *G* = 0 and *G* = 1, respectively.

¶¶ Note that a similar trend was experimentally detected, e.g., for the diffusion of doxorubicin^181^ drug molecules in confined silica nanoslits in Ref.^182^, while *deeper* minima of the velocity-ACFs at later time shifts were found, in contrast, for chromatin dynamics in computer simulations of viscoelastic subdiffusion of chromosomal loci in Ref.^183^.

*** Note that the direct comparison of the computed ACFs from simulations with those of, for instance, fractional Brownian motion with a *constant* Hurst exponent *H* is not straightforward due to the fact that the time-local anomalous-diffusion exponents in the simulations *vary* strongly with lag time, see Fig. 3.

## Notes

### Competing Interest Statement

The authors have declared no competing interest.

